# SARS-CoV-2 Omicron Spike recognition by plasma from individuals receiving BNT162b2 mRNA vaccination with a 16-weeks interval between doses

**DOI:** 10.1101/2021.12.21.473679

**Authors:** Debashree Chatterjee, Alexandra Tauzin, Lorie Marchitto, Shang Yu Gong, Marianne Boutin, Catherine Bourassa, Guillaume Beaudoin-Bussières, Yuxia Bo, Shilei Ding, Annemarie Laumaea, Dani Vézina, Josée Perreault, Laurie Gokool, Chantal Morrisseau, Pascale Arlotto, Éric Fournier, Aurélie Guilbault, Benjamin Delisle, Inès Levade, Guillaume Goyette, Gabrielle Gendron-Lepage, Halima Medjahed, Gaston De Serres, Cécile Tremblay, Valérie Martel-Laferrière, Daniel E. Kaufmann, Renée Bazin, Jérémie Prévost, Sandrine Moreira, Jonathan Richard, Marceline Côté, Andrés Finzi

## Abstract

Continuous emergence of SARS-CoV-2 variants of concern (VOC) is fueling the COVID-19 pandemic. Omicron (B.1.1.529), is rapidly spreading worldwide. The large number of mutations in its Spike raised concerns about a major antigenic drift that could significantly decrease vaccine efficacy and infection-induced immunity. A long interval between BNT162b2 mRNA doses was shown to elicit antibodies that efficiently recognize Spikes from different VOCs. Here we evaluated the recognition of Omicron Spike by plasma from a cohort of SARS-CoV-2 naïve and previously-infected individuals that received their BNT162b2 mRNA vaccine 16-weeks apart. Omicron Spike was recognized less efficiently than D614G, Alpha, Beta, Gamma and Delta Spikes. We compared to plasma activity from participants receiving a short (4-weeks) interval regimen. Plasma from individuals of the long interval cohort recognized and neutralized better the Omicron Spike compared to those that received a short interval. Whether this difference confers any clinical benefit against Omicron remains unknown.

## INTRODUCTION

SARS-CoV-2 variants are constantly evolving under immune selective pressure. Ongoing mutational events in the viral genome leads to the emergence of variants with unique properties including increased transmission capabilities and resistance to antibodies elicited by both natural infection and vaccination. Based on transmission capabilities, virulence, and vaccine effectiveness, SARS-CoV-2 variants are classified as variants of concern (VOCs), variant of interest (VOIs) or variants under monitoring (VUMs)(WHO, 2021). In late 2020, the Alpha (B.1.1.7) variant emerged. The N501Y Spike mutation increased its affinity for the ACE2 receptor, leading to increased transmissibility (Davies et al., 2021; Prevost et al., 2021; Rambaut et al., 2020). The accumulation of E484K and K417N/T mutations along with N501Y in the receptor binding domain (RBD) led to the emergence of Beta (B.1.351) and Gamma (P.1) lineages, which rapidly spread worldwide (Amanat et al., 2021; ECDC, 2020; Tang et al., 2021). In April 2021, the Delta (B.1.617.2) variant emerged and quickly spread to most countries (Allen et al., 2022; Planas et al., 2021b), but is rapidly being replaced by Omicron (B.1.1.529). The World Health Organization designated Omicron as a VOC on November 26, 2021(WHO, 2021). Omicron accumulated more than 30 mutations in its Spike, raising concerns about a major antigenic drift that could significantly decrease vaccine efficacy.

Here we evaluated the recognition of the Omicron Spike by plasma from a cohort of SARS-CoV-2 naïve and previously-infected individuals that received the two BNT162b2 mRNA vaccine doses 16-weeks apart. We compared these responses to those elicited in individuals receiving a short dose interval regimen (4-weeks). Plasma from vaccinated previously-infected individuals recognized more efficiently all tested Spikes (D614G, Alpha, Beta, Gamma, Delta and Omicron) than those from naïve vaccinated individuals. Omicron Spike was recognized less efficiently than D614G, Alpha, Beta, Gamma and Delta Spikes. However, plasma from individuals receiving a long interval recognized and neutralized better the Omicron Spike compared to those that received a short interval.

## RESULTS

### Recognition of Spike variants by plasma from vaccinated individuals

The antigenic profile of D614G, Alpha, Beta, Gamma, Delta and Omicron Spikes was assessed with plasma collected three weeks (V3) and 4 months (V4) after the second dose of the BNT162b2 mRNA vaccine administered with a 16-weeks interval between doses (Figure 1A) (Tauzin et al., 2021a). Briefly, 293T cells were transfected with plasmids coding for full-length Spike variants. Two days post-transfection, cells were incubated with the indicated plasmas followed by flow cytometry analysis, as described (Anand et al., 2021; Beaudoin-Bussieres et al., 2020; Gasser et al., 2021; Prevost et al., 2020; Tauzin et al., 2021a; Tauzin et al., 2021b). VOCs Spike expression levels were normalized to the signal obtained with the conformationally independent anti-S2 neutralizing CV3-25 antibody (Li et al., 2021; Prevost et al., 2021; Ullah et al., 2021) that efficiently recognized and neutralized all VOCs Spike, including Omicron (Supplemental Figure 1). Using plasma from previously-infected individuals or from naïve double vaccinated individuals we observed a significant increase of recognition of all tested Spikes upon vaccination (Figure 1B-G), in agreement with previous observations (Stamatatos et al., 2021; Tauzin et al., 2021a; Tauzin et al., 2021b). In all cases, the Omicron Spike was significantly less recognized than all other Spikes, with the exception of the Beta variant (Figure 1D, H-J). At V3 and V4, levels of plasma binding against Omicron Spike in previously-infected individuals were similar to those against Delta Spike in naïve individuals (Figure 1 I and J). In agreement with previous observations (Tauzin et al., 2021a), Spike recognition declined more rapidly in the naïve group compared to previously-infected individuals.

**Figure 1.**
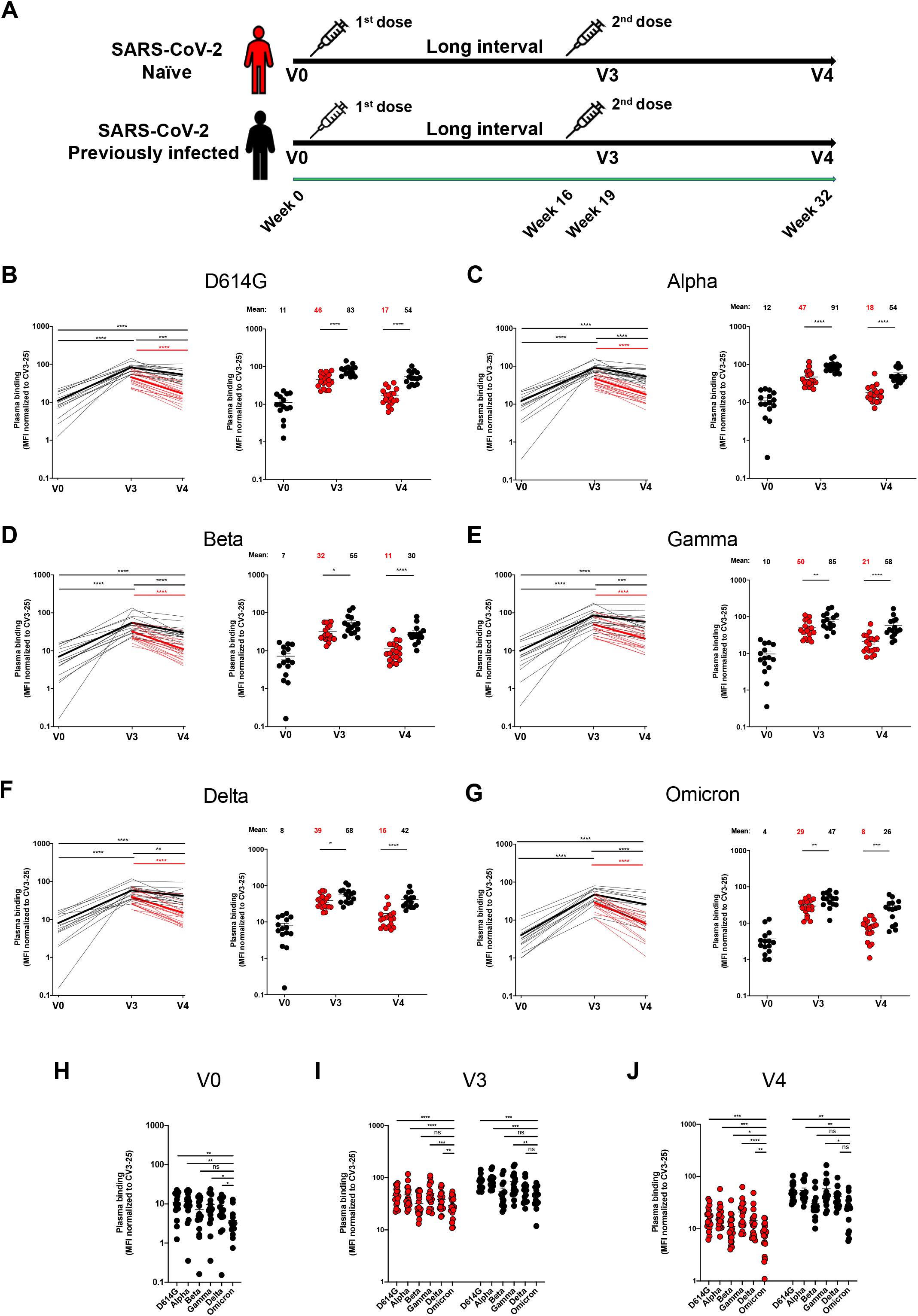
Binding of vaccine-elicited antibodies to SARS-CoV-2 Spike variants. (**A**) SARS-CoV-2 vaccine cohort design. (**B-G**) 293T cells were transfected with the indicated full-length Spike from different SARS-CoV-2 variants (D614G, Alpha, Beta, Gamma, Delta and Omicron) and stained with the CV3-25 Ab or with plasma collected 3 weeks (V3) or 4 months (V4) after a second dose administered with a 16 week interval. Samples were analyzed by flow cytometry. The values represent the median fluorescence intensities (MFI) normalized by CV3-25 Ab binding and presented as percentages of CV3-25 binding (**B-G, Left panels**). Each curve represents the normalized MFIs obtained with the plasma of one donor at every time point. The mean of each group is represented by a bold line. (**Right panels**) Plasma samples were grouped in different time points (V0, V3 and V4). (**H-J**) Comparison of Spike recognition by plasma from naïve and previously-infected donors, represented by red and black points respectively. Error bars indicate means ± SEM. For naïve donors, n=20 at V3 and V4. For previously infected donors vaccinated with two doses, n=15 at V0, V3 and V4. Statistical significance was tested using (**B-G Left panels, H, I, J**) a Wilcoxon test or (**B-G Right panels**) a Mann-Whitney test. (* P < 0.05; ** P < 0.01; *** P < 0.001; **** P < 0.0001; ns, non-significant)

### Impact of the interval between mRNA vaccine doses on Omicron Spike recognition and neutralization

Recent reports suggested that vaccine regimens with a delayed boost elicits stronger humoral responses than the approved, short interval, regimen (Grunau et al., 2021; Tauzin et al., 2021a). The long regimen interval has been associated with good vaccine efficacy against different VOCs (Skowronski et al., 2021). We therefore compared the capacity of plasma from naïve vaccinated individuals that received the second dose with an interval of sixteen weeks (median [range]: 111 days [76–120 days]) to those obtained from 19 SARS-CoV-2 naïve donors who received their two doses four weeks apart (median [range]: 29 days [22–34 days]) (Table 1 and Figure 2A). As shown in Figure 2B, plasma from naïve individuals that received a 16-weeks interval between the two doses, recognized significantly better all tested Spike, including Omicron, than plasma from individuals who received a short interval between doses (4 weeks). The 16-weeks interval regimen elicited significantly better neutralization activity against pseudoviral particles bearing the D614G, Beta, Delta and Omicron Spike (Figure 2C). Strikingly, this increased neutralization was more pronounced for the Omicron Spike (8.9 fold increase) comparatively to the others emerging variant Spikes (D614G, Beta and Delta) (2.2 - 4.2 fold increase). This suggests that the delayed boosting in naïve individuals facilitates antibody maturation resulting in enhanced breadth able to provide detectable levels of recognition and neutralization against Omicron.

**Table 1.** Characteristics of the vaccinated SARS-CoV-2 cohorts.

|  |  | SARS-CoV-2 Naïve |  | SARS-CoV-2 Previously infected |
| --- | --- | --- | --- | --- |
|  |  | Two doses<br>Short interval<br>(n=19) | Two doses<br>Long interval<br>(n=25) | Two doses<br>Long interval<br>(n=15) |
| Age |  | 39 (20-74) | 50 (21-62) | 47 (29-65) |
| Sex | Male (n) | 12 | 11 | 10 |
|  | Female (n) | 7 | 14 | 5 |
| Days between symptom onset and V0<sup>a</sup> |  | N/A | N/A | 191 (85-234) |
| Days between symptom onset and the 1<sup>st</sup> dose<sup>a</sup> |  | N/A | N/A | 274 (166-321) |
| Days between the 1<sup>st</sup> and 2<sup>nd</sup> dose<sup>a</sup> |  | 29 (22-34) | 111 (76-120) | 110 (90-134) |
| Days between the 2<sup>nd</sup> dose and V3<sup>a</sup> |  | 22 (12-53) | 21 (14-34) | 22 (13-51) |
| Days between the 2<sup>nd</sup> dose and V4<sup>a</sup> |  | N/A | 112 (103-125) | 113 (90-127) |

<sup>a</sup> Values displayed are medians, with ranges in parentheses.

**Figure 2.**
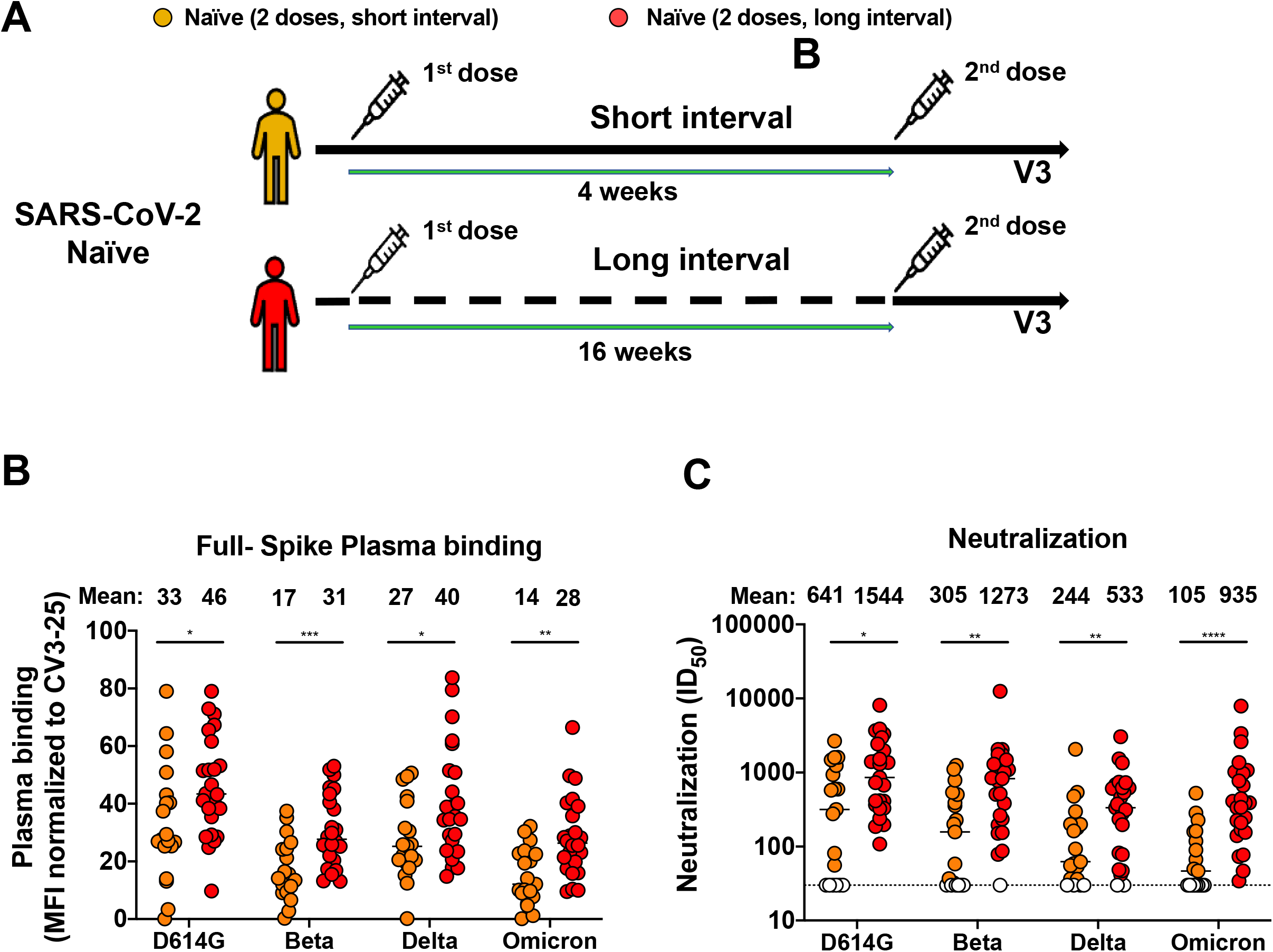
Omicron Spike recognition and neutralization with plasma from naïve individuals that received a short versus a long mRNA vaccine dose interval. (**A**) SARS-CoV-2 vaccine cohort design. (**B**) 293T cells were transfected with the full-length Spike from different SARS-CoV-2 variants (D614G, Beta, Delta and Omicron) and stained with the CV3-25 Ab or with plasma from naïve donors who received a short (4 weeks, yellow) or long (16 weeks, red) interval between doses collected three weeks after the second dose (V3) and analyzed by flow cytometry. The values represent the MFI normalized by CV3-25 Ab binding and presented as percentages of CV3-25 binding. (**C**) Neutralizing activity was measured by incubating pseudoviruses bearing indicated SARS-CoV-2 Spikes (D614G, Beta, Delta and Omicron), with serial dilutions of plasma for 1 h at 37°C before infecting 293T-ACE2 cells. Neutralization half maximal inhibitory serum dilution (ID_50_) values were determined using a normalized non-linear regression using GraphPad Prism software. Undetectable measures are represented as white symbols, and limits of detection are plotted. Error bars indicate means ± SEM. (**** P < 0.0001). For naïve donors vaccinated with the short interval, n=19. For naïve donors vaccinated with the long interval, n=25 Statistical significance was tested using a Mann-Whitney test (* P < 0.05; ** P < 0.01; *** P < 0.001;**** P < 0.0001).

## DISCUSSION

In the province of Québec, Canada, like in other jurisdictions worldwide, the prevalence of Omicron increased dramatically from the first case detected on November 23 to being the dominant variant less than one month later. Compared to the reference Wuhan-Hu-1 strain, the Omicron variant carries over 50 non-synonymous mutations within its genome, more than 30 of which are located in the gene coding for the Spike glycoprotein. Several of these mutations affecting the RBD, the N-terminal domain (NTD) and the furin cleavage domain were observed in other VOCs (Viana R., 2021), which is consistent with positive selection of favorable mutations. Previous *in vitro* studies already showed the association of some of these mutations with increased infectivity, ACE2 interaction (N501Y, P681H) (Gong et al., 2021; Saito et al., 2021) or immune evasion (K417N, N440K, G446S, S477N, E484A/K, Q493R) (Baum et al., 2020; Clark et al., 2021; Greaney et al., 2021a; Greaney et al., 2021b; Greaney et al., 2021c; Liu et al., 2020; Rappazzo et al., 2021; Starr et al., 2021; Weisblum et al., 2020). This unprecedented accumulation of Spike mutations raised concern about a major antigenic drift that could significantly decrease the efficacy of current vaccines (Andrews et al., 2021; Khoury et al., 2021; Schmidt et al., 2021b).

To get a better understanding of the antigenic profile, we compared the antigenicity of the Omicron Spike to those from D614G, Alpha, Beta, Gamma and Delta VOCs. We used plasma from naïve and previously-infected individuals who received their two doses of the BNT162b2 mRNA vaccine 16 weeks apart. In agreement with previous observations, we found that previously-infected vaccinated individuals recognized more efficiently all Spikes than naïve individuals at the two timepoints analyzed (3 weeks and 4 months post second dose) (Stamatatos et al., 2021; Tauzin et al., 2021a; Tauzin et al., 2021b). Interestingly, we observed that recognition of all Spikes, including Omicron, decreased more rapidly in naïve than previously-infected individuals, as reported (Tauzin et al., 2021a). The three antigenic exposures (infection + 2 doses) of previously-infected individuals compared to the two exposures in double-vaccinated naïve individuals possibly explain their more sustained humoral response, suggesting that an additional exposition to the Spike antigen in the form of a third vaccine dose could elicit similar responses. Alternatively, if infection elicits a qualitatively broader humoral response linked to epitopes located outside the Spike glycoprotein, the effect of a third dose in naïve individuals may remain qualitatively different. Independently of their infection history, all plasma recognized significantly less efficiently the Omicron Spike compared to Spikes from other VOCs (Figure 1).

Since recent studies have shown that recognition of full-length Spikes at the surface of transfected 293T cells strongly correlates with recognition of primary airway epithelial cells infected with authentic viruses as well as with antibody-dependent cellular cytotoxicity (Ding et al., 2022), these results suggest that Fc-mediated effector functions against Omicron could also be affected. Of note, low Spike recognition translated into increased Omicron neutralization resistance (Figure 2C). In agreement with previous observations (Stamatatos et al., 2021; Tauzin et al., 2021a; Tauzin et al., 2021b), plasma from vaccinated previously-infected individuals recognized more efficiently Omicron and all other VOCs than vaccinated naïve individuals (Figure 1). As naïve double-vaccinated individuals have been well protected against the delta variant, the observation of similar levels of plasma binding against Delta Spike in naïve individuals and those against Omicron Spike in previously-infected individuals may be important. This suggests that the benefits of hybrid immunity also apply to Omicron but this hypothesis will need confirmation through vaccine effectiveness studies.

Several reports (Cele et al., 2021; Garcia-Beltran et al., 2021; Planas et al., 2021a; Schmidt et al., 2021a; Zhang et al., 2021) have shown neutralization resistance using plasma from naïve donors who received the approved regimen of the BNT162b2 mRNA vaccine (3-4 weeks vaccine interval). Strikingly, we observed that plasma from naïve vaccinated donors who received their two doses according to the approved short three-four-week interval, recognized and neutralized significantly less efficiently Omicron compared to the long (16-weeks interval). For all individuals, the level of Omicron Spike recognition remains dramatically lower than for the ancestral Spike, the antigen used in the current vaccines and these levels decrease over time. Therefore, it will be important to determine in epidemiological studies if the vaccine interval advantage, as measured by these *in vitro* parameters, confers any clinical benefit against Omicron.

## ACKNOWLEDGMENTS

The authors are grateful to the donors who participated in this study. The authors thank the CRCHUM BSL3 and Flow Cytometry Platforms for technical assistance. We thank Dr. Stefan Pöhlmann (Georg-August University, Germany) for the plasmid coding for SARS-CoV-2 Spike Wuhan-Hu-1 strain. We also thank Amélie Boivin and Yves Grégoire at Héma-Québec for helping to access the samples from the PLASCOV Biobank and all the plasma donors who participate in this biobank. This work was supported by le Ministère de I’Économie et de I’Innovation du Québec, Programme de soutien aux organismes de recherche et d’innovation to A.F. and by the Fondation du CHUM. This work was also supported by a Canadian Institutes of Health Research (CIHR) foundation grant #352417, by a CIHR operating Pandemic and Health Emergencies Research grant #177958, a CIHR stream 1 and 2 for SARS-CoV-2 Variant Research to A.F., and by an Exceptional Fund COVID-19 from the Canada Foundation for Innovation (CFI) #41027 to A.F. and D.E.K. Work on variants presented was also supported by the Sentinelle COVID Quebec network led by the LSPQ in collaboration with Fonds de Recherche du Québec Santé (FRQS) to A.F. This work was also partially supported by a CIHR COVID-19 rapid response grant (OV3 170632) and CIHR stream 1 SARS-CoV-2 Variant Research to MC. A.F. is the recipient of Canada Research Chair on Retroviral Entry no. RCHS0235 950-232424. MC is a Tier II Canada Research Chair in Molecular Virology and Antiviral Therapeutics. V.M.L. is supported by a FRQS Junior 1 salary award. C,T. is the recipient of the Pfizer/Université de Montréal Chair on HIV translational research. D.E.K. is a FRQS Merit Research Scholar. G.B.B. is the recipient of a FRQS PhD fellowship and J.P. is the recipient of a CIHR PhD fellowship. A.L. was supported by MITACS Accélération postdoctoral fellowships. The funders had no role in study design, data collection and analysis, decision to publish, or preparation of the manuscript. We declare no competing interests.

## AUTHOR CONTRIBUTIONS

D.C., J.R., S.M., M.C., and A.F. conceived the study. D.C., A.T., J.R., M.C. and A.F. designed experimental approaches. D.C., A.T., L.M., S.Y.G., M.B., C.B., G.B.-B., Y.B., S.D., A.L., D.V., E.F., A.G., B.D., I.L., J.Prévost, J.R. and A.F. performed, analyzed, and interpreted the experiments. J.Perreault, L.G., C.M., P.A., C.T., V.M.-L., D.E.K. and R.B. collected and provided clinical samples. G.D.S., C.T., V.M.-L., and D.E.K. provided scientific input related to VOCs and vaccine efficacy. D.C., J.R., and A.F. wrote the manuscript with inputs from others. Every author has read, edited, and approved the final manuscript.

## DECLARATION OF INTERESTS

A.F. has filed a provisional patent application on the anti-Spike monoclonal antibody CV3-25.

## STAR METHODS

### RESOURCE AVAILABILITY

#### Lead contact

Further information and requests for resources and reagents should be directed to and will be fulfilled by the lead contact, Andrés Finzi.

#### Materials availability

All unique reagents generated during this study are available from the Lead contact without restriction.

#### Data and code availability

- All data reported in this paper will be shared by the lead contact upon request.
- This paper does not report original code.
- Any additional information required to reanalyze the data reported in this paper is available from the lead contact upon request.

### EXPERIMENTAL MODEL AND SUBJECT DETAILS

#### Ethics Statement

All work was conducted in accordance with the Declaration of Helsinki in terms of informed consent and approval by an appropriate institutional board. Blood samples were obtained from donors who consented to participate in this research project at CHUM (19.381) and from plasma donors who consented to participate in the Plasma Donor Biobank at Hema-Quebec (PLASCOV; REB-B-6-002-2021-003). Plasma was isolated by centrifugation, and samples stored at −80°C and in liquid nitrogen, respectively, until use.

#### Human subjects

The study was conducted in 25 SARS-CoV-2 naïve individuals (11 males and 14 females; age range: 21-62 years) vaccinated with a long interval, 19 SARS-CoV-2 naïve individuals (12 males and 7 females; age range: 20-74 years) vaccinated with a short interval and 15 SARS-CoV-2 previously-infected individuals (10 males and 5 females; age range: 29-65 years) vaccinated with a long interval. All this information is summarized in Table 1. No specific criteria such as number of patients (sample size), gender, clinical or demographic were used for inclusion, beyond PCR confirmed SARS-CoV-2 infection in adults.

#### Plasma and antibodies

Plasma from SARS-CoV-2 naïve and previously-infected donors were collected, heat-inactivated for 1 hour at 56°C and stored at −80°C until ready to use in subsequent experiments. The conformationally independent S2-specific monoclonal antibody CV3-25 (Gong et al., 2021; Jennewein et al., 2021; Li et al., 2021; Prevost et al., 2021; Ullah et al., 2021) was used as a positive control and to normalize Spike expression in our flow cytometry assays, as described (Gong et al., 2021; Tauzin et al., 2021a; Tauzin et al., 2021b). Alexa Fluor-647-conjugated goat anti-human Abs (Invitrogen) were used as secondary antibodies to detect plasma binding in flow cytometry experiments.

#### Cell lines

293T human embryonic kidney cells (obtained from ATCC) were maintained at 37°C under 5% CO_2_ in Dulbecco’s modified Eagle’s medium (DMEM) (Wisent) containing 5% fetal bovine serum (FBS) (VWR) and 100 μg/ml of penicillin-streptomycin (Wisent). The 293T-ACE2 cell line was previously reported (Prévost et al., 2020).

### METHOD DETAILS

#### Plasmids

The plasmids encoding the SARS-CoV-2 Spike variants; D614G, B.1.1.7, B.1.351, P.1 and B.1.617.2 were previously described (Beaudoin-Bussieres et al., 2020; Gong et al., 2021; Li et al., 2021; Tauzin et al., 2021a; Tauzin et al., 2021b). The plasmids encoding the B.1.1.529 Spike was generated by overlapping PCR using a codon-optimized wild-type SARS-CoV-2 Spike gene (GeneArt, ThermoFisher) that was synthesized (Biobasic) and cloned in pCAGGS as a template. The B.1.1.529 Spike coding sequence was derived from the sequence ID EPI_ISL_6640919. This sequence initially contained the Q493K substitution, as previously reported (Cameroni et al., 2021; Schmidt et al., 2021a; Shah and Woo, 2021). The ECDC (European Centre for Disease Prevention and Control) later informed that Omicron spike actually have an R mutation at position 493. We therefore generated and used an Omicron Spike bearing the Q493R mutation for the full manuscript (Figure 1–2 and Supplemental Figure 1). Nevertheless, we compared whether the nature of the residue (either K or R) at this position impacted plasma recognition and/or neutralization activity; no significant differences were observed (Supplemental Figure 2).

#### Cell surface staining and flow cytometry analysis

293T were transfected with full-length SARS-CoV-2 Spikes and a green fluorescent protein (GFP) expressor (pIRES2-eGFP; Clontech) using the calcium-phosphate method. Two days post-transfection, Spike-expressing 293T cells were stained with the CV3-25 Ab (5 μg/mL) as control or plasma from vaccinated individuals (1:250 dilution) for 45 min at 37°C. AlexaFluor-647-conjugated goat anti-human IgG (1/1000 dilution) were used as secondary Abs. The percentage of Spike-expressing cells (GFP+ cells) was determined by gating the living cell population based on viability dye staining (Aqua Vivid, Invitrogen). Samples were acquired on a LSR II cytometer (BD Biosciences), and data analysis was performed using FlowJo v10.7.1 (Tree Star). The conformationally-independent anti-S2 antibody CV3-25 was used to normalize Spike expression, as reported (Gong et al., 2021; Li et al., 2021; Prevost et al., 2021; Ullah et al., 2021). CV3-25 was shown to be effective against all Spike variants (Gong et al., 2021; Li et al., 2021; Prevost et al., 2021; Ullah et al., 2021) and (Supplemental figure 1). The Median Fluorescence intensities (MFI) obtained with plasma were normalized to the MFI obtained with CV3-25 (Gong et al., 2021; Li et al., 2021; Prevost et al., 2021; Ullah et al., 2021) and presented as percentage of CV3-25 binding.

#### Virus neutralization assay

To produce SARS-CoV-2 pseudoviruses, 293T cells were transfected with the lentiviral vector pNL4.3 R-E-Luc (NIH AIDS Reagent Program) and a plasmid encoding for the indicated S glycoprotein (D614G, Alpha, Beta, Gamma, Delta or Omicron) at a ratio of 10:1. Two days post-transfection, cell supernatants were harvested and stored at −80°C until use. For the neutralization assay, 293T-ACE2 target cells were seeded at a density of 1×10^4^ cells/well in 96-well luminometer-compatible tissue culture plates (Perkin Elmer) 24h before infection. Pseudoviral particles were incubated with several plasma dilutions (1/50; 1/250; 1/1250; 1/6250; 1/31250) for 1h at 37°C and were then added to the target cells followed by incubation for 48h at 37°C. For CV3-25 neutralization, pseudoviral particles were incubated with increasing concentrations of CV3-25 (0.01, 0.0316, 0.1, 0.316, 1 and 3.16 μg/mL) for 1h at 37°C and were then added to the target cells followed by incubation for 48h at 37°C. Cells were lysed by the addition of 30 μL of passive lysis buffer (Promega) followed by one freeze-thaw cycle. An LB942 TriStar luminometer (Berthold Technologies) was used to measure the luciferase activity of each well after the addition of 100 μL of luciferin buffer (15mM MgSO4, 15mM KH2PO4 [pH 7.8], 1mM ATP, and 1mM dithiothreitol) and 50 μL of 1mM d-luciferin potassium salt (Prolume). The neutralization half-maximal inhibitory dilution (ID_50_) represents the plasma dilution to inhibit 50% of the infection of 293T-ACE2 cells by pseudoviruses.

### QUANTIFICATION AND STATISTICAL ANALYSIS

#### Statistical analysis

Symbols represent biologically independent samples from SARS-CoV-2 naïve or PI individuals. Lines connect data from the same donor. Statistics were analyzed using GraphPad Prism version 8.0.1 (GraphPad, San Diego, CA). Every dataset was tested for statistical normality and this information was used to apply the appropriate (parametric or nonparametric) statistical test. P values <0.05 were considered significant; significance values are indicated as * P<0.05, ** P<0.01, *** P<0.001, **** P<0.0001, ns, non-significant.

---

**Supplemental Figure 1.**
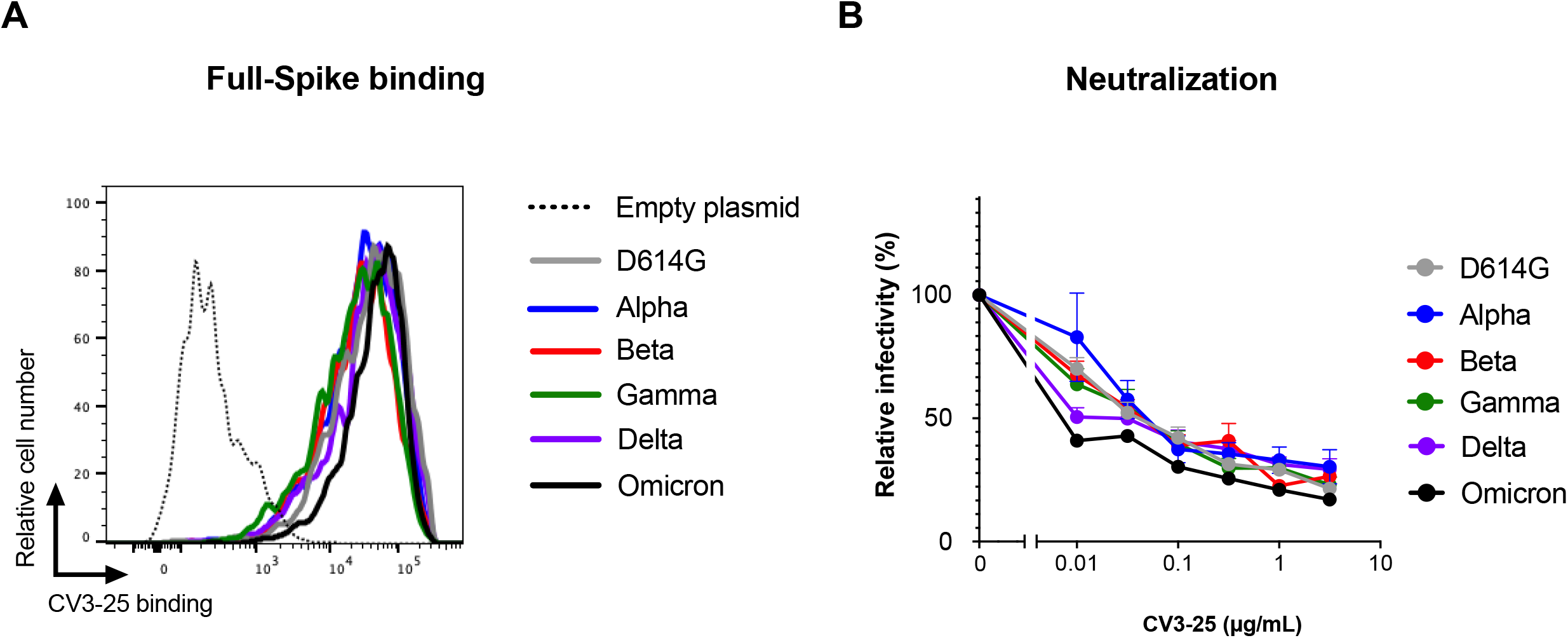
Recognition and neutralization of different VOCs Spikes by the anti-S2 neutralizing CV3-25 antibody. (A) 293T cells were transfected with the full-length Spikes from different VOCs (D614G, Alpha, Beta, Gamma, Delta and Omicron), stained with the CV3-25 Ab and analyzed by flow cytometry. Shown are histograms showing a representative staining on Spike-expressing GFP+ cells. (B) CV3-25 neutralizing activity against pseudoviral particles bearing the different VOC Spikes.

**Supplemental Figure 2.**
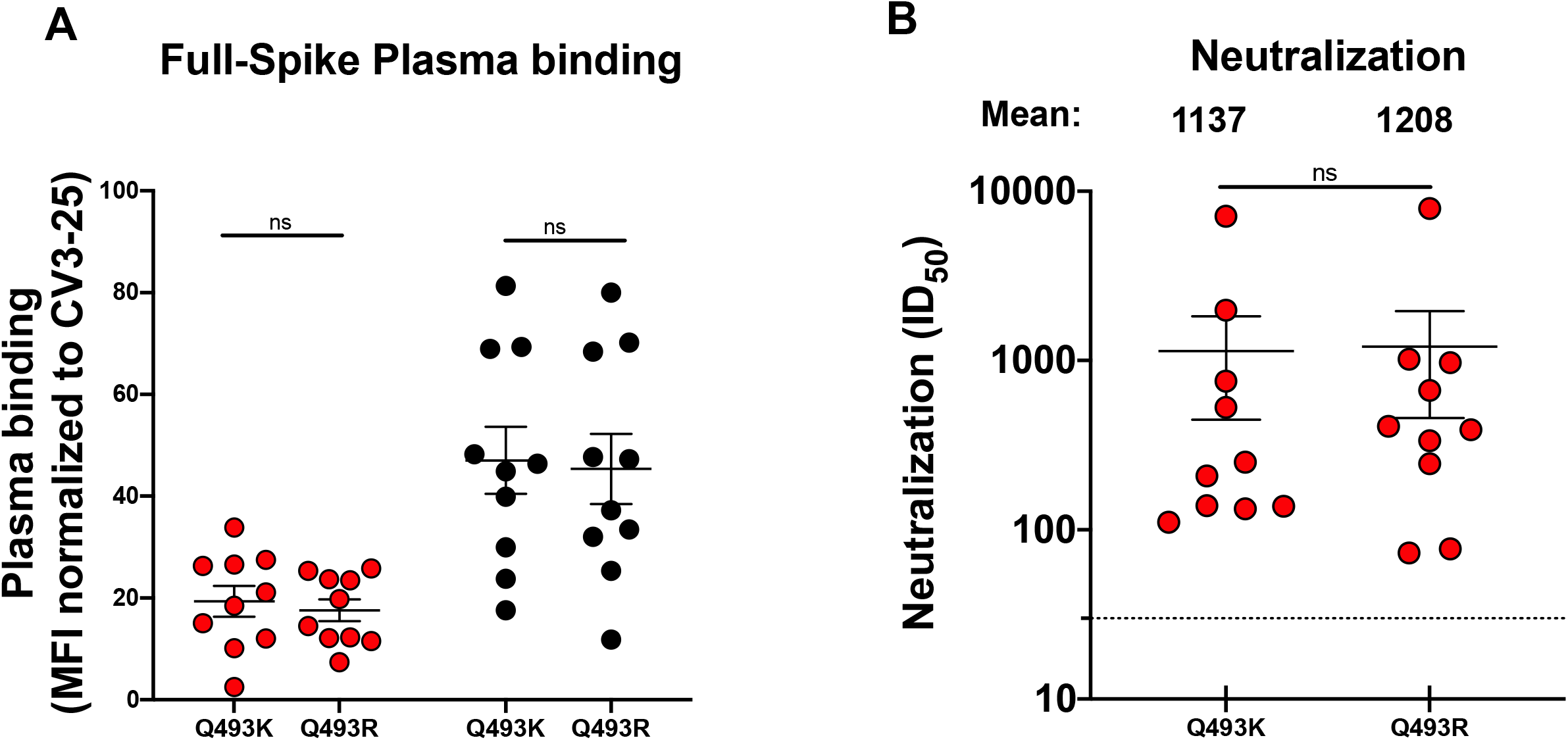
Recognition and neutralization of Omicron Spikes with Q493K or Q493R changes. (A) 293T cells were transfected with the full-length Spikes of Omicron possessing either Q493K or Q493R mutation, stained with the CV3-25 Ab or with plasma collected 3 weeks (V3) after the second dose with a 16-week interval from naïve or previously-infected donors, represented by red and black points respectively and analyzed by flow cytometry. (B) Neutralizing activity was measured by incubating pseudoviruses bearing SARS-CoV-2 Spikes of the two different Omicron mutants as mentioned above, with serial dilutions of plasma collected 3 weeks (V3) after the second dose with a 16-week interval from naïve donors for 1 h at 37°C before infecting 293T-ACE2 cells. Neutralization half maximal inhibitory serum dilution (ID_50_) values were determined using a normalized non-linear regression using GraphPad Prism software. Error bars indicate means ± SEM. Statistical significance was tested using a Wilcoxon test (ns, non-significant).

